# Complement Component 3 expressed by the endometrial ectopic tissue is involved in the endometriotic lesion formation through mast cell activation

**DOI:** 10.1101/2020.11.19.389536

**Authors:** C. Agostinis, S. Zorzet, A. Balduit, G. Zito, A. Mangogna, Paolo Macor, F. Romano, M. Toffoli, B. Belmonte, A. Martorana, V. Borelli, G. Ricci, U. Kishore, R. Bulla

**Affiliations:** Institute for Maternal and Child Health, IRCCS Burlo Garofolo, 34137, Trieste, Italy; Department of Life Sciences, University of Trieste, 34127, Trieste, Italy; Tumor Immunology Unit, Human Pathology Section, Department of Health Sciences, University of Palermo, Palermo, Italy; Department of Health Promotion, Mother and Child Care, Internal Medicine and Medical Specialties, University of Palermo, Palermo, Italy; Department of Medical, Surgical and Health Science, University of Trieste, Trieste, Italy; Biosciences, College of Health, Medicine and Life Sciences, Brunel University London, London, United Kingdom

## Abstract

The pathophysiology of endometriosis (EM) is an excellent example of immune dysfunction, reminiscent of tumor microenvironment as well. Here, we report that an interplay between C3 and mast cells (MCs) is involved in the pathogenesis of ectopic EM. C3 is at the epicenter of the regulatory feed forward loop, amplifying the inflammatory microenvironment, in which the MCs are protagonists. Thus, C3 can be considered a marker of EM and its local synthesis can promote the engraftment of the endometriotic cysts. We generated a murine model of EM via injection of minced uterine tissue from a donor mouse, into the peritoneum of the recipient mice. The wild type mice showed greater amount of cyst formation in the peritoneum compared to C3 knock-out mice. This study offers an opportunity for novel therapeutic intervention in EM, a difficult to treat gynecological condition.

**Summary:** C3 produced by the endometriotic tissue is involved in the lesion development through mast cell activation

## Introduction

Endometriosis (EM) is a chronic condition that affects about 5-10% of women in reproductive age. EM is characterized by pain and infertility as a consequence of the presence of functional endometrial tissue outside the uterine cavity (*1*). The most common locations for the ectopic implants are the ovaries, peritoneum, and the utero-sacral ligaments. The presence of ectopic tissue in these areas induces a condition of chronic inflammation. Current evidence suggests that immune dysfunction is the most likely causative factor for the EM pathogenesis. In particular, the pathways involved in immune-cell recruitment, cell-adhesion, and inflammatory processes encourage the implantation and survival of endometriotic lesions. Recently, it has been shown that the complement system is one of the most predominant pathways altered in EM (*2*).

The complement system is an important part of the innate immunity and acts as a bridge between innate and acquired immune system. It is involved in host defense against infectious agents and altered self. Three different molecules are responsible for the recognition phase of complement: C1q, Mannose-Binding Lectin (MBL), and C3. The binding of the recognition molecules to the target ligands initiate the three different complement pathways: the classical (via C1q), alternative (via C3) and lectin (via MBL) pathway (*3, 4*). The three pathways eventually converge in the formation of the C3 and C5 convertases. This then results in the generation of the main effector molecules of the complement system: the opsonins C3b and C4b, the anaphylatoxins C3a, C4a and C5a, and the Membrane Attack Complex (MAC) that causes the target cell lysis. The small complement fragments, anaphylatoxins, cause local inflammatory responses by acting, for instance, on mast cells (MCs), inducing an increase in blood flow, vascular permeability, and leukocyte recruitment (*5, 6*).

The current consensus is that EM involves a local pelvic inflammatory process with altered function of immune-related cells in the peritoneal environment. Recent studies suggest that the peritoneal fluid (PF) of women with EM contains an increased number of activated macrophages that secrete various humoral mediators locally, including growth factors, cytokines and possibly, oxidative products (*7*), leading to the development and progression of EM and EM-associated infertility. Although the contributions of specific immune cell subsets and their mediators to the onset and the course of the inflammatory process in endometrial lesions are still poorly understood, evidence suggests that MCs are crucially involved in the inflammatory process associated with EM. High numbers of degranulated MCs have been found in endometriotic lesions (*8–13*).

Complement component C3 is expressed by the epithelial cells found in endometriotic tissue (*14–16*). In this study, we examined differential expression of C3 in eutopic and ectopic tissues derived from EM lesions and its consequences in the pathogenesis of EM using human EM tissues and C3 gene-deficient mice.

## Results

### C3 is present in ectopic, but not in eutopic, endometrium and is locally expressed by endometriotic cells

We initially confirmed the presence of C3 in human endometriotic tissue sections by immunofluorescence (IF, fig. S1) and by immunohistochemical assays (Fig. 1), in uterine and ectopic endometrium. IHC, using anti-human C3 polyclonal antibody, showed moderate cytoplasmic expression of C3 by endometrial stromal cells, and rarely in some glandular epithelial cells in proliferative as well as secretory endometrium (Fig.1, A and B). A strong signal was present in adnexal and peritoneal EM (Fig. 1, C to G). C3 was found to be widely distributed in the EM tissue, with variable intensity; it was mostly localized in the glandular-like structures and in the vessels.

**Fig. 1.**
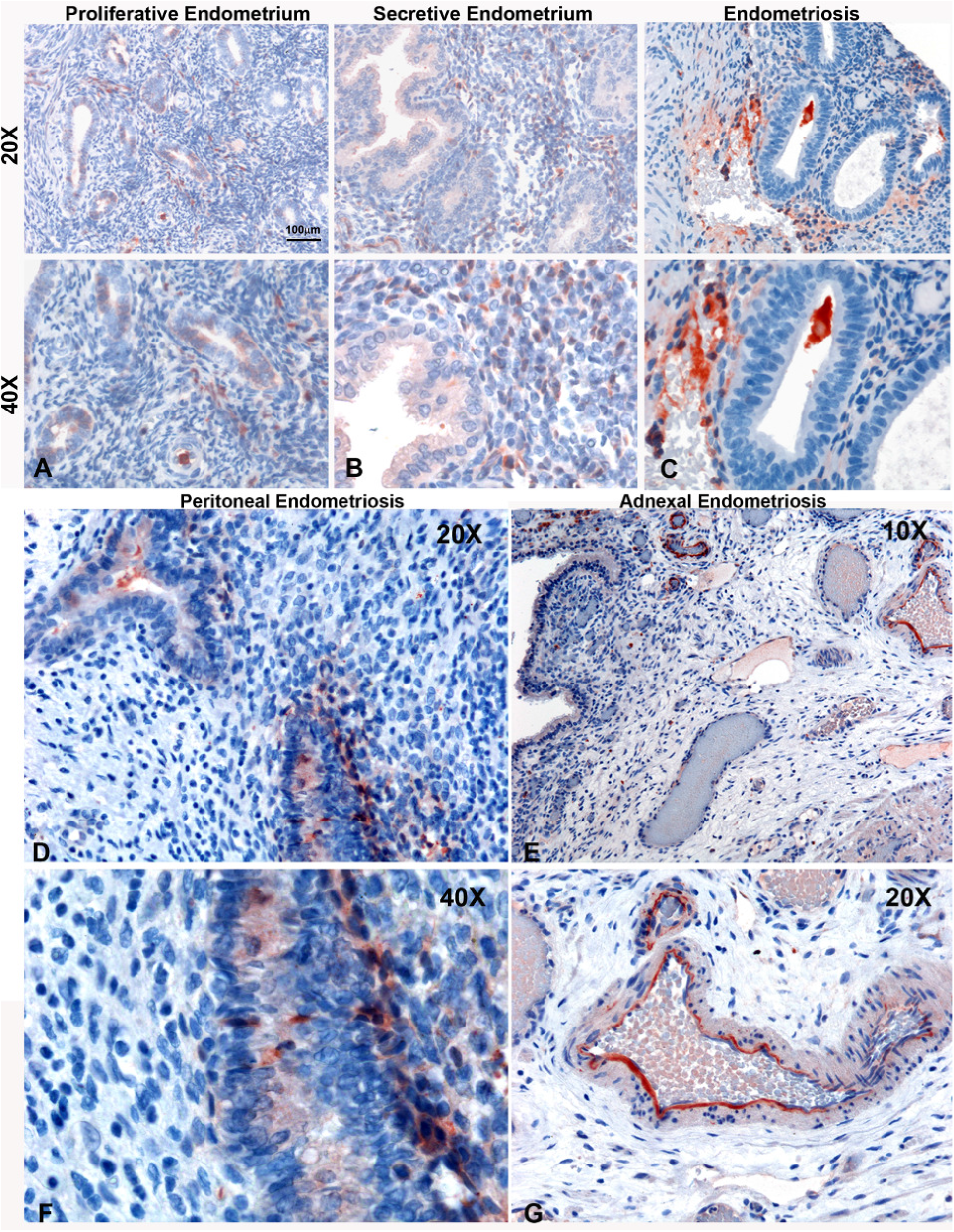
C3 expression by human eutopic and ectopic endometrial tissue. Representative microphotographs showing expression of C3 by IHC in proliferative normal endometrium (A), secretive normal endometrium (B), and endometriotic tissue sections (C). In the lower panel: C3 IHC on abdominal wall (D and F) and adnexal (E and F) endometriosis (EM). AEC (red) chromogen was used to visualize the binding of anti-human C3 antibodies. Nuclei were counterstained in blue with Harris Hematoxylin; scale bars, 100μm.

In order to investigate the local synthesis of C3, total RNA was isolated from EM cysts and C3 gene expression was analyzed by RT-qPCR. Ectopic EM tissues expressed high levels of C3 transcripts compared to the eutopic normal endometrium (Fig. 2A). To assess the contribution of EM epithelial cells to the local C3 production, we isolated EM epithelial/stromal cells from human EM cysts, which showed the characteristic epithelial phenotype (Fig. 2B); ~98% of the cells were positive for the epithelial marker, Cytokeratin 8/18 (fig. S2). We analyzed their expression of C3 by RT-qPCR using HepG2 (hepatocyte cell line) as a calibration control. AN3CA cells were used to assess the expression of C3 by normal epithelial endometrial cells. Because the IHC analysis revealed a marked positivity in the EM vessels, the expression of C3 was also investigated in endometriotic endothelial cells (EEC) and corresponding eutopic cells (uterine microvascular endothelial cells; UtMEC). We observed a very low expression of C3 transcript by both endothelial cell types (fig. S3). Interestingly, EM epithelial/stromal cells expressed a higher amount of C3 compared to AN3CA and HepG2 cells (Fig. 2C). In addition, these cells secreted the C3 protein in the culture supernatant, as measured via ELISA (Fig. 2D). To establish the specificity and exclusivity of C3 expression, we also measured C5 mRNA level; we could not detect C5 transcripts either in the EM tissue or cultured EM epithelial cells (data not shown).

**Fig. 2.**
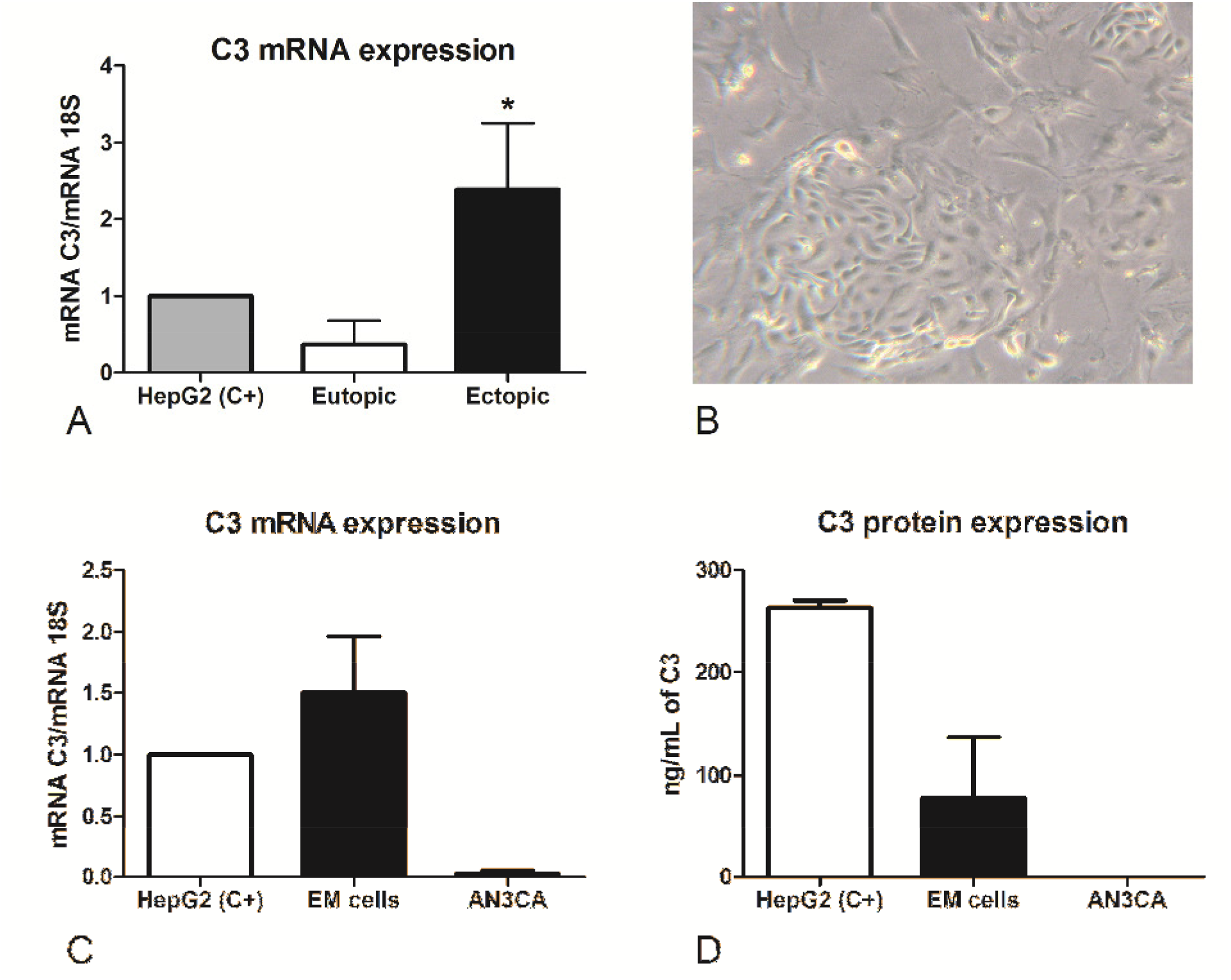
C3 is locally produced in EM tissue. (A) The gene expression of C3 by endometriotic cysts (n = 4) was investigated by RT-qPCR and compared to normal uterus (n= 3) using HepG2 hepatocyte cells as calibrator (AU = 1). 18S was used as the housekeeping gene. Data are expressed as mean ± standard error. * p<0.05 (Mann-Whitney U Test). (B) Example of morphological features of endometrial cells isolated from EM cyst biopsies. Images were acquired by phase-contrast microscope, Leica original magnification: 200×. (C) The C3 mRNA expression was measured by RT-qPCR in EM cells (n= 4) and compared to an endometrial cell line, AN3CA. Data are expressed as mean ± standard error. * p<0.05 (Mann-Whitney U Test). (D) Protein levels of C3 were assessed by ELISA in the supernatant of cells maintained in culture for 60 h. Data are expressed as mean ± standard error.

### Pro-inflammatory stimuli induce C3 expression by normal endometrial cells, in an in vitro model of EM

Because the ectopic endometrial microenvironment is characterized by inflammation, we set up an in vitro model of EM, stimulating the endometrial cell line, AN3CA, with pro-inflammatory cytokines. Our results demonstrated that normal endometrial cells under resting conditions were unable to produce C3; however, when stimulated with TNF-α, and to a lesser extent with IL-1β, cells started to express C3 mRNA (Fig. 3A), and secrete an increased level of C3 protein, as measured by ELISA (Fig. 3B) as well as by western blot analysis (Fig. 3, C and D). We demonstrated also that the stimulation of C3 transcription induced by TNF-α was dose-dependent (fig. S4).

**Fig. 3.**
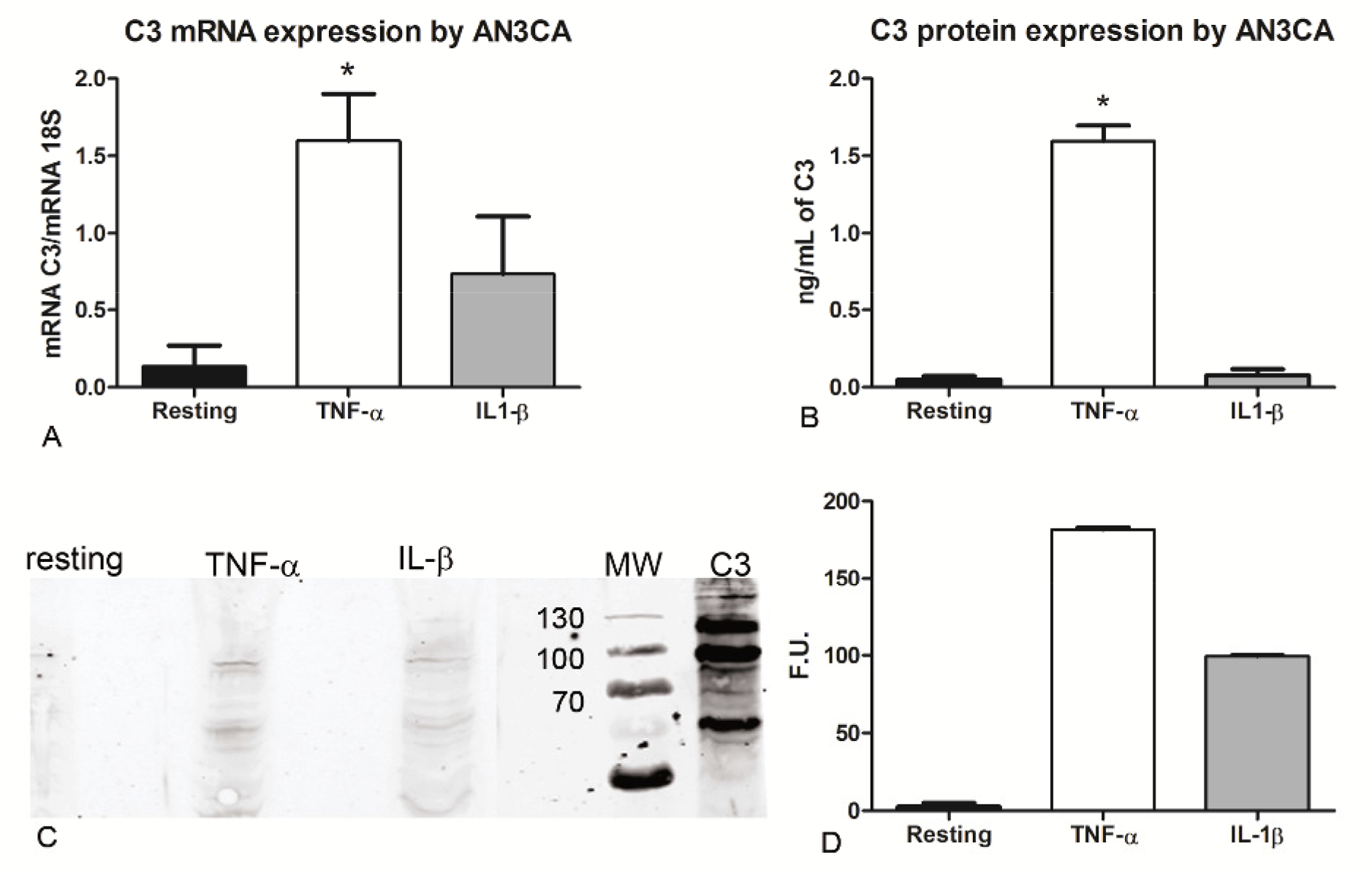
C3 is up-regulated by pro-inflammatory cytokines. The stimulation of AN3CA cells with pro-inflammatory cytokines induced the up-regulation of C3 expression. (A) AN3CA cells were ON stimulated with TNF-α (100ng/mL) or IL-1β (5ng/mL) and the C3 expression was analyzed by RT-qPCR. Data are expressed as mean of three independent experiments conducted in double ± standard error. * p<0.05 (Wilcoxon matched pairs test). (B) Measurement of protein level of C3 by ELISA in AN3CA cell culture supernatant stimulated for 36h with TNF-α (100ng/mL) or IL-1 β (5ng/mL). (C and D) The expression of C3 was evaluated in cell lysates by western blot analysis and the intensity of the bands was measured with Odyssey-LICOR scanner. Data are expressed as mean of three independent experiments conducted in double ± standard error. * p<0.05.

### C3^-/-^ mice are refractory to developing EM cysts in a syngeneic in vivo model of EM

To further investigate the role of C3 in EM pathogenesis, we set up a syngeneic model of EM in C57BL/6 WT or C3 gene-deficient (C3^-/-^) mice. Estrus was induced in donor animals via administration of estradiol. Then, the minced uterus of the donor mouse was injected in the peritoneum of 2 recipient mice (Fig. 4A). After 3 weeks, the animals were sacrificed, and the peritoneal cysts were counted. WT mice developed a higher number of cysts compared to C3 deficient mice (Fig. 4B), indicating that lack of C3 could prevent the EM cyst formation.

**Fig. 4.**
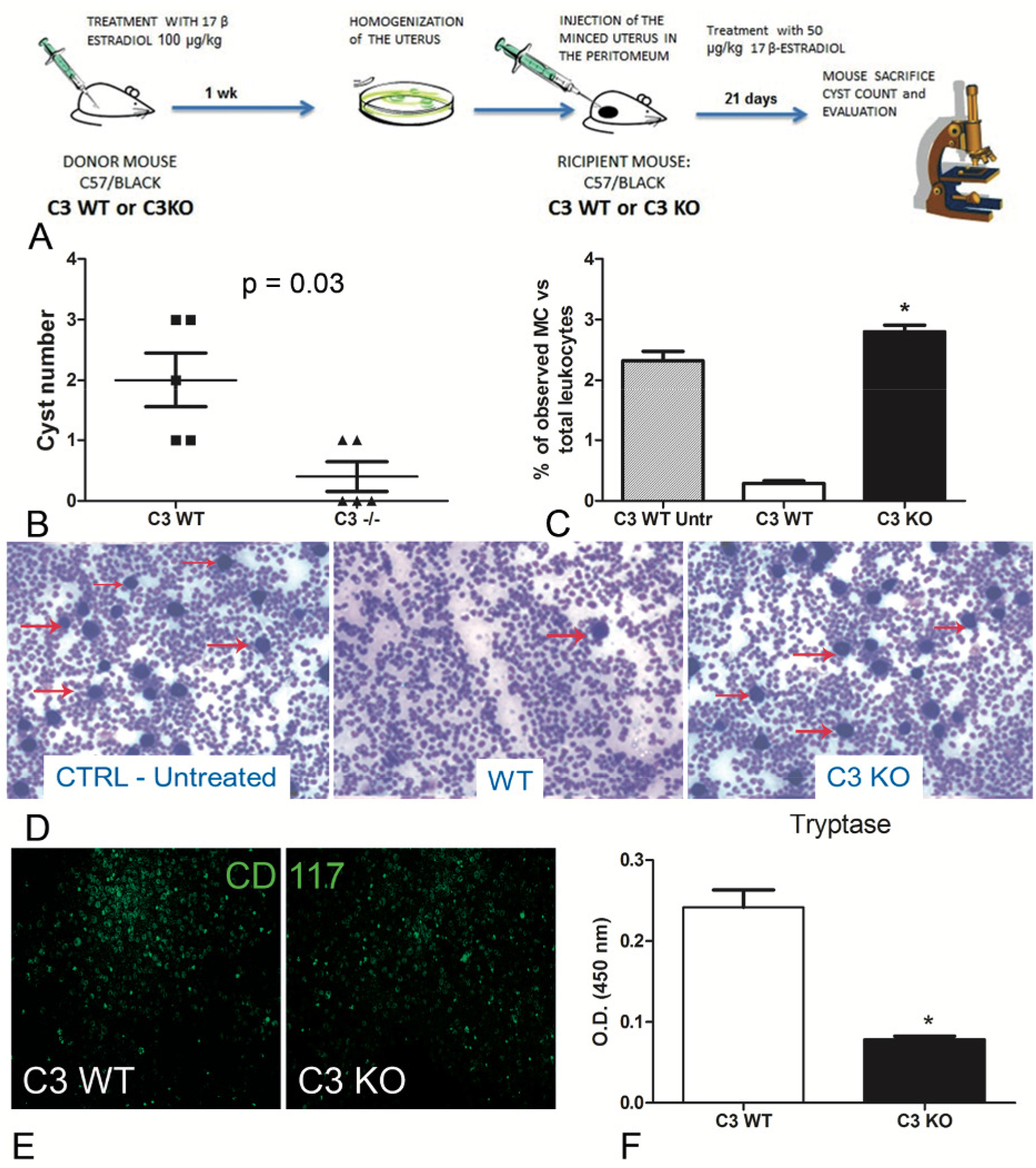
In vivo syngeneic mouse model of EM. (A) Treatment regimen of C3 WT and gene-deficient mice for generating EM in vivo model. Five C3-/- and WT mice each were injected (i.p.) with minced uterus of a donor mouse; after 3 weeks, the peritoneal cyst formation was evaluated. (B) Number of cysts counted in wild-type (WT, *n* = 5) or C3-/- (*n* = 5) mice. Mann-Whitney test *P* = 0.03. (C and D) Representative images of cytocentrifuged peritoneal washing of untreated WT, EM-induced WT and C3-/-mice (respectively), stained with Giemsa and counted with ImageJ software (Particle Analysis Tool) to obtain relative percentage between total leukocytes and mast cell (MC)/basophil number. MCs/basophils are identified as blue big dots indicated by red arrows. Original magnification 100×. (E) Representative images of cytocentrifuged peritoneal lavage of EM-induced WT mice stained with FITC conjugated anti-mouse CD117. Original magnification 100×. (F) Biochemical characterization of tryptase enzyme present in peritoneal lavage of WT vs C3-/-mice by ELISA. * p<0.05.

### Peritoneal washing isolated from WT mice with EM present more degranulated MCs compared to C3^-/-^ mice

We then analyzed the peritoneal washing of WT and C3^-/-^ mice for the presence of infiltrating leukocytes. The samples were cytocentrifuged and the cells were stained with Giemsa. A high number of leukocytes was observed in the peritoneal washing of both WT and C3^-/-^ treated mice. The number of infiltrating leukocytes was comparable with those observed in WT untreated (CTRL Untr) mice. Surprisingly, the fluids collected from the WT animals contained a lower number of MCs, compared to the C3^-/-^ mice (Fig. 4, C and D). Next, we stained the cytocentrifuged leucocytes with a monoclonal antibody to CD117 (a marker of mature MCs). The peritoneal washing collected from the WT mice contained CD117 positive cells as well (Fig. 4E). We wanted to confirm if the MCs present in WT peritoneal washing were degranulated. We thus measured the levels of MC enzyme, tryptase, in the murine peritoneal washing (Fig. 4F). The ELISA for tryptase demonstrated that WT mice with EM had a higher amount tryptase compared to C3^-/-^ or CTRL mice.

### C3a level is higher in the PF of EM patients, which can act on EM MCs

C3a is one of the most important stimuli for MC activation. Thus, we measured the levels of the C3a in the PF of EM patients, and compared them with those obtained from myoma and fibroma patients (women undergoing laparoscopy, without alterations of peritoneal cavity environment). Our results showed that the PF of EM patients had significantly higher level of C3a compared to the control patients (Fig. 5A).

**Fig. 5.**
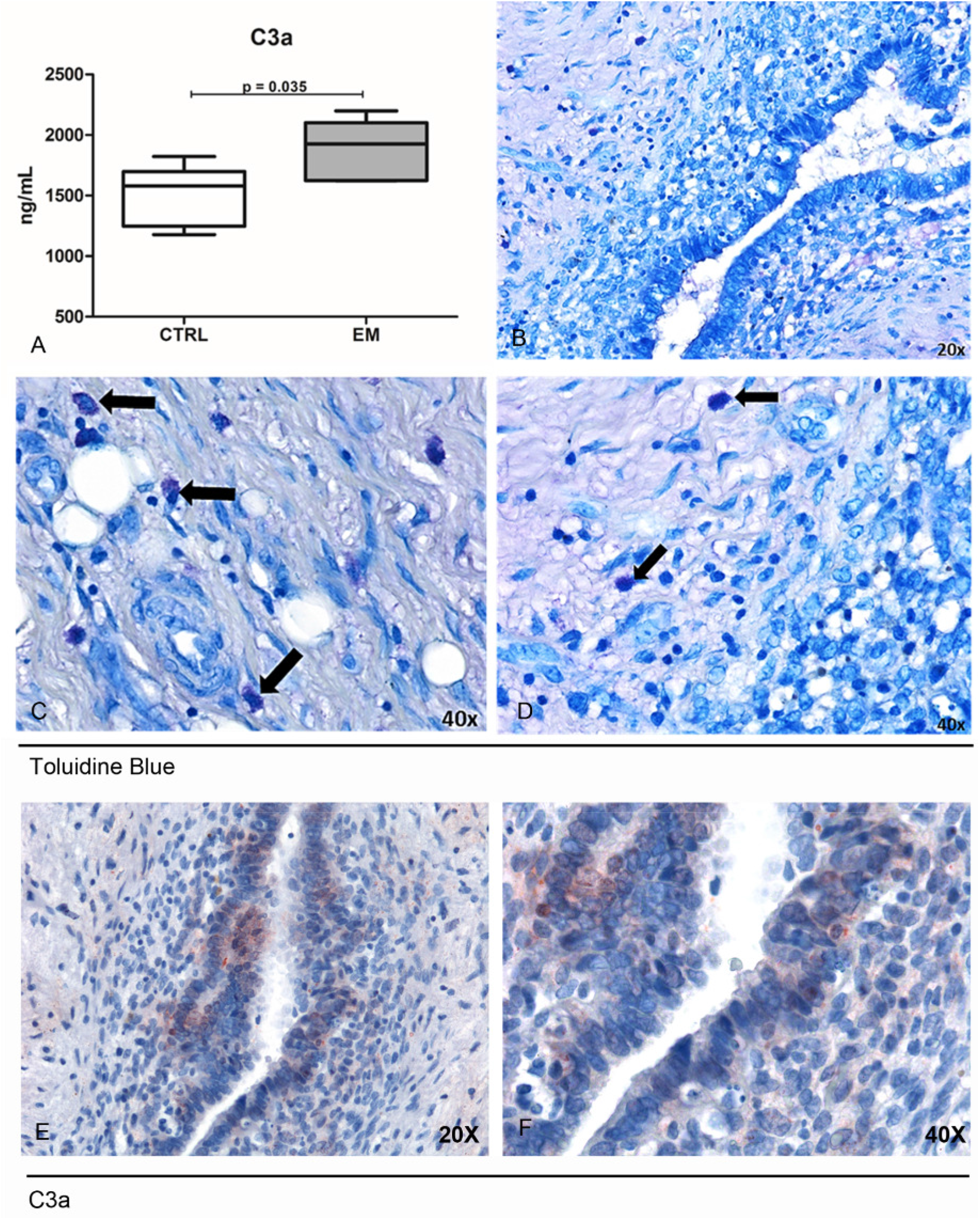
Peritoneal Fluid (PF) derived from EM patients presented elevated levels of C3a that likely acts on MCs of the EM tissue. (A) C3a ELISA evaluation of PF isolated from EM patients (n = 7) compared to control patient group (n = 6). (B to D) Toluidine blue staining of human EM tissue sections for the evaluation of MCs presence. Black arrows indicated MCs. (E and F) Immunohistochemical analysis of C3a in EM tissue sections. AEC (red) chromogen was used to visualize the binding of anti-human C3a antibodies.

We next considered that the C3a present in the peritoneal cavity of EM patients could stimulate the MCs within the EM lesions. We therefore stained human EM tissue sections with toluidine blue to highlight MCs present in the tissue. The histochemical analysis confirmed that EM lesions were rich in MCs (Fig. 5, B to D), compared to normal endometrium (fig. S5); the IHC for C3a on the same sections corroborated the presence of this anaphylatoxin as well (Fig. 5, E and F).

### C3a is involved in an auto-amplifying loop of inflammation between MCs and EM cells

In order to reconstruct the cross-talk between EM cells and MCs, we set up a co-culture assay, seeding endometrial cell line (AN3CA) at the bottom of a 24-well plate and placing the HMC-1 (a MC cell line) onto the upper chamber of a transwell system (1 μm diameter pores). To understand the role of C3a in this cross-talk, the co-culture was also performed in the presence of a pool of EM-PF and/or a blocking anti-C3a antibody. AN3CA cells, cultured with HMC-1 without any other stimulus, expressed low levels of C3. When HMC-1 cells were stimulated with EM-PF in the co-culture, AN3CA cells began to express higher quantities of C3 (Fig. 6A). Surprisingly, AN3CA cells, stimulated with only EM-PF (in absence of HMC-1), were not able to express C3; they produced C3 only in response to EM-PF stimulated HMC-1, or to TNF-α (used as a positive control). Pre-incubation of EM-PF with a blocking monoclonal anti-C3a antibody completely abrogated the effect of EM-PF stimulation, bringing it to resting values (Fig. 6B).

**Fig. 6.**
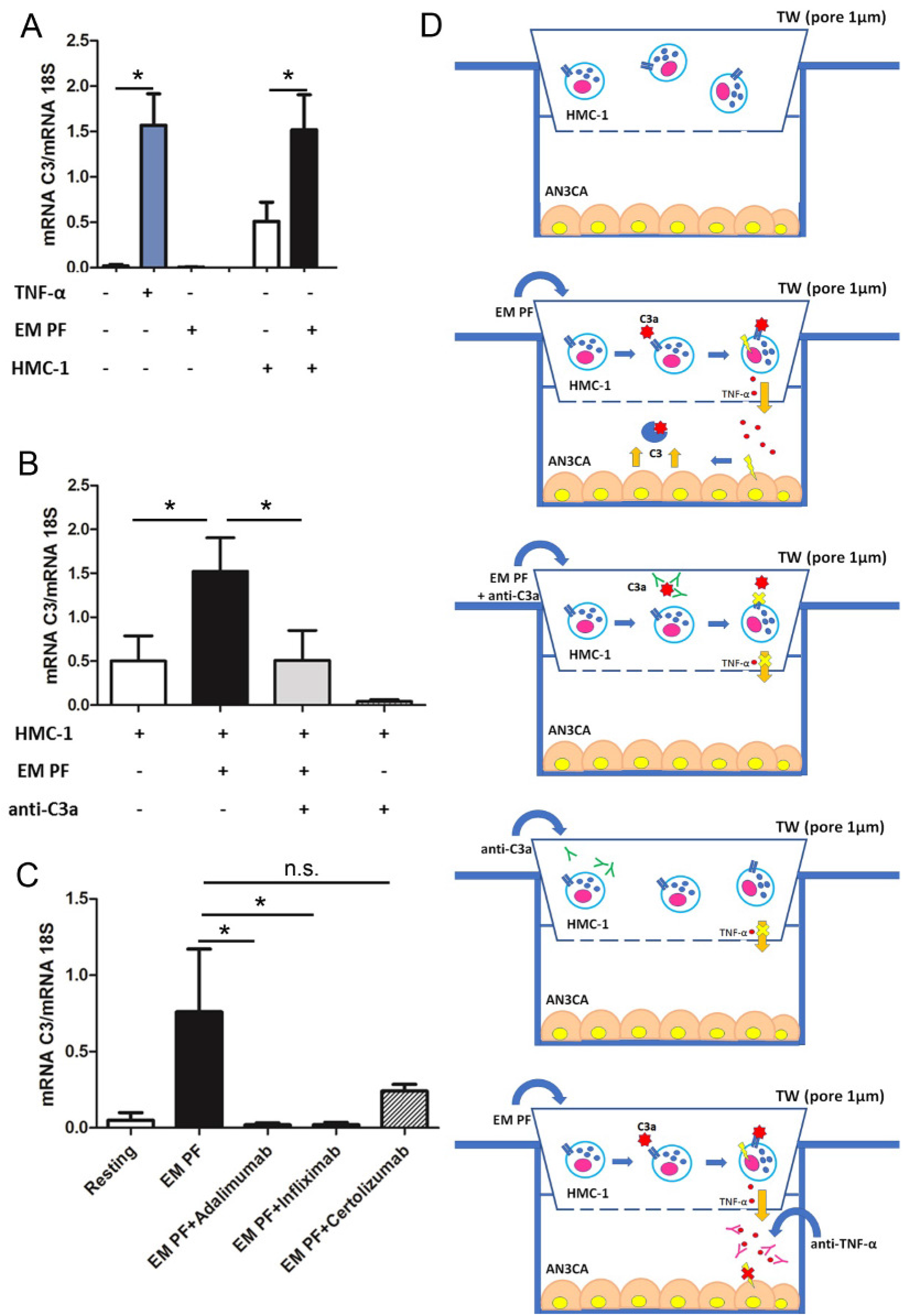
The co-culture of MCs with endometrial cells, in the presence of EM-PF, induced the expression of C3, which was inhibited by C3a blocking antibody. (A) The C3 gene expression was evaluated by RT-qPCR on endometrial AN3CA cells alone (Resting), stimulated with TNF-α (+TNF-α) or with a pool of EM peritoneal fluid (+EM-PF); or co-cultured with MCs alone (+HMC-1), or stimulated with PFA as positive control (+HMC-1+PFA) or with EM-PF (+HMC-1+ EM PF). Similar experiment was performed in the presence of anti-C3a blocking antibody (B), and the explained by the graphical abstract in (C). (D) Graphical representation of the blocking co-culture experiments. Data are expressed as mean of three independent experiments conducted in double ± standard error. * p<0.05.

With a view to gauge a potential novel immunotherapeutic approach for EM, we investigated in our co-culture model the effect of anti-TNF-α antibodies. Infliximab and Adalimumab (both in clinical use) showed a strong effect in blocking C3 expression by AN3CA, whereas Certolizumab did not block significantly the effects induced by the EM-PF on HMC-1 cells (Fig. 6C).

## Discussion

There is sufficient evidence for the expression of complement components and regulators in human endometrium (*17–20*). Dysregulation of the complement system is largely considered to play an important role in the pathogenesis of EM (*2, 21, 22*). The importance of the complement proteins in EM was highlighted recently by high-throughput studies (*2, 21–23*). The complement system appears to dominate the chronic inflammation in EM and remains a precipitating factor in EM-associated ovarian cancer. Complement factors C3, C4A, C7, factor D, factor B, factor H, and mannose-associated serine protease 1 (MASP1), are differentially expressed in EM compared to normal uterine tissues (*2*). Furthermore, single-nucleotide polymorphisms in the C3 gene show association with an increased risk of EM and EM-associated infertility (*15*).

The presence of C3 in EM tissue was first highlighted in 1980 (*24*). Subsequent studies confirmed the presence of higher levels of C3 in the EM lesions (*2, 14–16, 21, 23, 25–28*). However, how C3 contributes to the pathogenic mechanisms in EM has not been elucidated yet. We show, in this study, the presence of C3 in eutopic and ectopic endometrium (Fig. 1); a strong signal was observed in EM tissue (Fig 1, C to F). C3 was found to be widely distributed in the EM lesions with variable intensity, being mostly localized in the glandular-like structures and in the vessels.

C3 appeared to be produced locally and in higher amounts in the ectopic tissues compared to the normal uterus (Fig. 2A). EM cells, isolated from ovarian cysts of women of fertile age, produced C3 in culture (Fig. 2, C and D) (*22, 26, 27*). We also sought to investigate the factors produced by endometrial implants located in the abdominal cavity, which influenced the C3 production, and consequent inflammatory microenvironment in the uterine endometrium (*25*).

A potential candidate for the upregulation of C3 expression by endometrial cells is the PF rich in pro-inflammatory factors (*29*). PF TNF-α concentrations are elevated in EM patients; its higher concentration correlates with the severity and stage of the disease (*30–32*). Furthermore, TNF-α and IL-1β are known to be elevated in EM milieu (*29, 33*). The stimulation of endometrial cells with TNF-α and IL-1β induced the production of C3. Surprisingly, the stimulation of endometrial stromal cells with EM-PF did not cause increased C3 gene expression, indicating that the concentration of TNF-α in the PF (around 50 pg/ml) was not sufficient for a direct endometrial cell activation (*31*). We wondered what the source of higher concentration of TNF-α in the endometrial tissue could be. A partial explanation came from the study of C3^-/-^ mice. We observed that WT mice developed a higher number of cysts and presented degranulated MCs in PF, as demonstrated by the presence of a considerably higher amount of tryptase, compared to C3^/-^ mice (Fig. 3). Thus, endometrial cells under pro-inflammatory stimuli express C3; the pro-inflammatory milieu can induce the generation of C3a, which in turn, can activate the MCs. Activated MCs can release histamine, in addition to secreting TNF-α (and a host of inflammatory factors).

One of the possible causes of complement activation in the EM microenvironment is the setting off of the coagulation cascade, which is caused by the typical periodic bleeding in the EM tissue. Thrombin cleaves C3 to C3a and C3b; activated platelets are also involved in C3 cleavage (*34*). Another activator of the C3 is heme that is released from hemoglobin during hemolysis; heme induces deposition of C3 fragments on the erythrocytes (*35*). Alternative pathway activation can occur through its up-regulator, properdin, binding to activated platelets promoting C3(H_2_O) recruitment and complement activation (*36*). In addition, stimulation of endothelial cells by C3a or other factors promptly induces expression of P-selectin, which by binding to C3b, induces the formation of C3 convertases (*37*).

We confirmed the presence of higher levels of C3a in the EM-PF compared to control non-EM healthy women (Fig. 5A). The presence of complement component (C4) and complement activation products (C3c SC5b-9) in the PF and in the sera of EM women have been previously described (*38*). Significantly higher PF levels of C1q, MBL and C1-INH in women with EM compared to control group has also been reported (*39*).

C3a present in the EM-PF can act, through C3aR interaction, which is abundantly expressed on the MCs present in the EM tissue (*9, 25*). The involvement of MCs in the EM lesion formation was previously investigated. A recent study showed that numbers of MCs in total as well as activated MCs were clearly increased in the EM lesions in both animals and humans. Thus, the use of MC stabilizers and inhibitors may prove to be effective in treating EM and its associated pain (*40*). An increased presence of activated and degranulated MCs in deeply infiltrating EM and its close histological relationship with nerves strongly suggests that MCs contribute to the development of pain and hyperalgesia in EM, possibly by a direct effect on the nerve structures (*10*). A diffuse infiltration of numerous MCs was observed throughout the stromal lesions, which exhibited degranulation; scattered granules were also observed. In the eutopic endometrium and normal uterine serosa of the EM patients and the controls, MCs were rarely detected (*11*). We identified MCs in the EM cyst tissues (Fig. 5, B to D). Very few cells were localized in the endometrial stroma, which is characteristic of endometrial gland-like regions. A good proportion of MCs were localized around blood vessels and the interstitium with fibrosis, that is, the fibrotic interstitium of endometrial cysts. MCs thus are likely to be involved in the development and progression of EM. Localization of MCs suggests its particularly close relationship with fibrosis and adhesion (*41, 42*).

In conclusion, we demonstrated that the complement component C3 is locally synthesized by ectopic endometrial tissue, while the normal uterine endometrium does not express C3. Normal endometrial cells, under pro-inflammatory stimuli, produce C3. C3 gene-deficient mice present less endometriotic lesions in a syngeneic EM mouse model. C3a seems to recruit massive MCs in the EM lesions, which can have a damaging pathogenic consequence in the ectopic EM. C3 seems central to a regulatory feed forward loop, and is able to amplify the inflammatory microenvironment, in which the MCs are protagonists. This study opens up a new window for the identification of new targets for treating EM.

## Materials and Methods

### Patients

Patients and control women were enrolled at the Institute for Maternal and Child Health, IRCCS Burlo Garofolo, Trieste, Italy. The study group (EM) consisted of a total of 7 women, diagnosed with moderate/severe and minimal/mild EM, according to the revised criteria of the American Society for Reproductive Medicine ASRM (*43*).

### Primary cell isolation and culture

Endometriotic cells and endothelial microvascular uterine cells (UtMEC) were isolated, as described earlier (*44*).

### Murine model of EM

C3^-/-^ mice were kindly provided by Prof. Marina Botto, Centre for Complement and Inflammation Research, Department of Medicine, Imperial College, London (UK) and generated as described previously (*45*). The mouse model of EM was adapted from Somigliana and Mariani (*46, 47*).

### Gene expression analysis

RNA was extracted from cells using the kit supplied by Norgen Biotek Corp. (Aurogene, Rome, Italy) according to the manifacturer’s instructions and reverse transcripted to cDNA through SuperMix kit (Bioline). qPCR was carried out on a Rotor-Gene 6000 (Corbett, Qiagen, Milan, Italy) using SYBR™ Green PCR Master Mix (Applied Biosystems, Milan, Italy). Supplementary Table S1 shows the primers used for RT-qPCR. The melting curve was recorded between 55°C and 99°C with a hold every 2s. The relative amount of gene expression in each sample was determined by the Comparative Quantification (CQ) method supplied as a part of the Rotor Gene 1.7 software (Corbett Research) (*48*). The relative amount of each gene was normalized with 18S and expressed as arbitrary units (AU), considering 1 AU obtained from HepG2 cells used as a calibrator.

### Measurement of C3a

The levels of C3a in PF from EM patients and C3 in cell culture supernatant were evaluated by ELISA kits purchased from Quidel.

### Co-culture and cell stimulation

A confluent 24 well-plate of AN3CA cells was stimulated overnight (ON) with 100ng/mL of TNF-α, or 5ng/mL of IL-1 β (both from PeproTech, ListerFish, Milan, Italy), or 10% of pooled EM-PF. For the co-culture study, the cells were seeded at confluence, in the lower part of a 1μm pore TW system (Corning, Milan, Italy), whereas the HMC-1 (2×10^5^/TW) was present in the upper part, in the presence of 10% of pooled EM-PF, with or without blocking anti-human C3a (MyBiosource, Aurogene, Milan, Italy; 20μg/mL). Subsequently, the cells were lysed for RNA extraction, the supernatant recovered and stored at −80°C.

## General

We thank Prof. Marina Botto (Department of Medicine, Imperial College, London, UK) for providing C3^-/-^ mice; Daisy Bertoni for the contribution to the study, Ghergana Topouzova for help with patient enrolment, and Prof. Carlo Pucillo and Barbara Frossi (University of Udine, Udine, Italy) for providing HMC-1 cell line.

## Author contributions

C.A., S.Z., and R.B. designed research; C.A., A.B., V.B. and S.Z. performed research; P.M., G.Z., A.Mar. and F.R. contributed new reagents/analytic tools; B.B., M.T. and A.Man. analyzed data; C.A., G.R. and R.B. conceived the study; R.B. and G.R. supervised the study; C.A., S.Z., B.B. and A.B. supervised experiments; C.A., A.B., U.K., A.Man. and R.B. wrote the paper.

## References

1. M. J. Schleedoorn, W. L. Nelen, G. A. Dunselman, N. Vermeulen, G. EndoKey, Selection of key recommendations for the management of women with endometriosis by an international panel of patients and professionals. Hum Reprod 31, 1208–1218 (2016).

2. S. Sturyawanshi et al., Complement pathway is frequently altered in endometriosis and endometriosis-associated ovarian cancer. Clin Cancer Res 20, 6163–6174 (2014).

3. M. J. Walport, Complement. First of two parts. N Engl J Med 344, 1058–1066 (2001).

4. M. J. Walport, Complement. Second of two parts. N Engl J Med 344, 1140–1144 (2001).

5. M. Noris, G. Remuzzi, Overview of complement activation and regulation. Semin Nephrol 33, 479–492 (2013).

6. M. M. Markiewski, J. D. Lambris, The role of complement in inflammatory diseases from behind the scenes into the spotlight. Am J Pathol 171, 715–727 (2007).

7. A. Capobianco, P. Rovere-Querini, Endometriosis, a disease of the macrophage. Front Immunol 4, 9 (2013).

8. R. Paula, Jr. et al., The intricate role of mast cell proteases and the annexin A1-FPR1 system in abdominal wall endometriosis. J Mol Histol 46, 33–43 (2015).

9. D. Kirchhoff, S. Kaulfuss, U. Fuhrmann, M. Maurer, T. M. Zollner, Mast cells in endometriosis: guilty or innocent bystanders? Expert Opin Ther Targets 16, 237–241 (2012).

10. V. Anaf et al., Pain, mast cells, and nerves in peritoneal, ovarian, and deep infiltrating endometriosis. Fertil Steril 86, 1336–1343 (2006).

11. M. Sugamata, T. Ihara, I. Uchiide, Increase of activated mast cells in human endometriosis. Am J Reprod Immunol 53, 120–125 (2005).

12. D. Kempuraj et al., Increased numbers of activated mast cells in endometriosis lesions positive for corticotropin-releasing hormone and urocortin. Am J Reprod Immunol 52, 267–275 (2004).

13. R. Konno et al., Role of immunoreactions and mast cells in pathogenesis of human endometriosis--morphologic study and gene expression analysis. Hum Cell 16, 141–149 (2003).

14. P. G. Signorile, A. Baldi, Serum biomarker for diagnosis of endometriosis. J Cell Physiol 229, 1731–1735 (2014).

15. L. A. Ruiz et al., Single-nucleotide polymorphisms in the lysyl oxidase-like protein 4 and complement component 3 genes are associated with increased risk for endometriosis and endometriosis-associated infertility. Fertil Steril 96, 512–515 (2011).

16. D. Bartosik, I. Damjanov, R. R. Viscarello, J. A. Riley, Immunoproteins in the endometrium: clinical correlates of the presence of complement fractions C3 and C4. Am J Obstet Gynecol 156, 11–15 (1987).

17. R. A. Sayegh, X. J. Tao, J. T. Awwad, K. B. Isaacson, Localization of the expression of complement component 3 in the human endometrium by in situ hybridization. J Clin Endocrinol Metab 81, 1641–1649 (1996).

18. K. P. Murray et al., Expression of complement regulatory proteins-CD 35, CD 46, CD 55, and CD 59-in benign and malignant endometrial tissue. Gynecol Oncol 76, 176–182 (2000).

19. A. Franchi, J. Zaret, X. Zhang, S. Bocca, S. Oehninger, Expression of immunomodulatory genes, their protein products and specific ligands/receptors during the window of implantation in the human endometrium. Mol Hum Reprod 14, 413–421 (2008).

20. J. R. Sherwin et al., Identification of novel genes regulated by chorionic gonadotropin in baboon endometrium during the window of implantation. Endocrinology 148, 618–626 (2007).

21. S. H. Ahn et al., Immune-inflammation gene signatures in endometriosis patients. Fertil Steril 106, 1420–1431 e1427 (2016).

22. H. Kobayashi et al., The ferroimmunomodulatory role of ectopic endometriotic stromal cells in ovarian endometriosis. Fertil Steril 98, 415–422 e411–412 (2012).

23. K. Rekker et al., High-throughput mRNA sequencing of stromal cells from endometriomas and endometrium. Reproduction 154, 93–100 (2017).

24. J. C. Weed, P. C. Arquembourg, Endometriosis: can it produce an autoimmune response resulting in infertility? Clin Obstet Gynecol 23, 885–893 (1980).

25. P. Bischof, D. Planas-Basset, A. Meisser, A. Campana, Investigations on the cell type responsible for the endometrial secretion of complement component 3 (C3). Hum Reprod 9, 1652–1659 (1994).

26. K. B. Isaacson, Q. Xu, C. R. Lyttle, The effect of estradiol on the production and secretion of complement component 3 by the rat uterus and surgically induced endometriotic tissue. Fertil Steril 55, 395–402 (1991).

27. K. B. Isaacson, C. Coutifaris, C. R. Garcia, C. R. Lyttle, Production and secretion of complement component 3 by endometriotic tissue. J Clin Endocrinol Metab 69, 1003–1009 (1989).

28. I. Flores et al., Molecular profiling of experimental endometriosis identified gene expression patterns in common with human disease. Fertil Steril 87, 1180–1199 (2007).

29. A. Fassbender, R. O. Burney, D. F. O, T. D’Hooghe, L. Giudice, Update on Biomarkers for the Detection of Endometriosis. Biomed Res Int 2015, 130854 (2015).

30. M. A. Bedaiwy, T. Falcone, Peritoneal fluid environment in endometriosis. Clinicopathological implications. Minerva Ginecol 55, 333–345 (2003).

31. M. A. Bedaiwy et al., Prediction of endometriosis with serum and peritoneal fluid markers: a prospective controlled trial. Hum Reprod 17, 426–431 (2002).

32. J. Eisermann, M. J. Gast, J. Pineda, R. R. Odem, J. L. Collins, Tumor necrosis factor in peritoneal fluid of women undergoing laparoscopic surgery. Fertil Steril 50, 573–579 (1988).

33. K. E. May et al., Peripheral biomarkers of endometriosis: a systematic review. Hum Reprod Update 16, 651–674 (2010).

34. M. M. Markiewski, B. Nilsson, K. N. Ekdahl, T. E. Mollnes, J. D. Lambris, Complement and coagulation: strangers or partners in crime? Trends Immunol 28, 184–192 (2007).

35. A. W. Pawluczkowycz, M. A. Lindorfer, J. N. Waitumbi, R. P. Taylor, Hematin promotes complement alternative pathway-mediated deposition of C3 activation fragments on human erythrocytes: potential implications for the pathogenesis of anemia in malaria. J Immunol 179, 5543–5552 (2007).

36. K. Fromell et al., Assessment of the Role of C3(H_2_O) in the Alternative Pathway. Front Immunol 11, 530 (2020).

37. R. A. Harrison, The properdin pathway: an “alternative activation pathway” or a “critical amplification loop” for C3 and C5 activation? Semin Immunopathol 40, 15–35 (2018).

38. J. Kabut, Z. Kondera-Anasz, J. Sikora, A. Mielczarek-Palacz, Levels of complement components iC3b, C3c, C4, and SC5b-9 in peritoneal fluid and serum of infertile women with endometriosis. Fertil Steril 88, 1298–1303 (2007).

39. J. Sikora et al., The role of complement components C1q, MBL and C1 inhibitor in pathogenesis of endometriosis. Arch Gynecol Obstet 297, 1495–1501 (2018).

40. M. M. Binda, J. Donnez, M. M. Dolmans, Targeting mast cells: a new way to treat endometriosis. Expert Opin Ther Targets 21, 67–75 (2017).

41. H. Fujiwara et al., Localization of mast cells in endometrial cysts. Am J Reprod Immunol 51, 341–344 (2004).

42. S. Matsuzaki et al., Increased mast cell density in peritoneal endometriosis compared with eutopic endometrium with endometriosis. Am J Reprod Immunol 40, 291–294 (1998).

43. Revised American Society for Reproductive Medicine classification of endometriosis: 1996. Fertil Steril 67, 817–821 (1997).

44. C. Agostinis et al., The combination of N-acetyl cysteine, alpha-lipoic acid, and bromelain shows high anti-inflammatory properties in novel in vivo and in vitro models of endometriosis. Mediators Inflamm 2015, 918089 (2015).

45. M. R. Wessels et al., Studies of group B streptococcal infection in mice deficient in complement component C3 or C4 demonstrate an essential role for complement in both innate and acquired immunity. Proc Natl Acad Sci U S A 92, 11490–11494 (1995).

46. E. Somigliana et al., Endometrial ability to implant in ectopic sites can be prevented by interleukin-12 in a murine model of endometriosis. Hum Reprod 14, 2944–2950 (1999).

47. M. Mariani et al., The selective vitamin D receptor agonist, elocalcitol, reduces endometriosis development in a mouse model by inhibiting peritoneal inflammation. Hum Reprod 27, 2010–2019 (2012).

48. R. D. McCurdy, J. J. McGrath, A. Mackay-Sim, Validation of the comparative quantification method of real-time PCR analysis and a cautionary tale of housekeeping gene selection. Gene Therapy & Molecular Biology 12, 15–24 (2008).

