## Supplemental Material for "Complement Component 3 expressed by the endometrial ectopic tissue is involved in the endometriotic lesion formation through mast cell activation"

**Title**

**Short Title**

- Role of complement component 3 in endometriosis

**Authors**

C. Agostinis[1], S. Zorzet[2], A. Balduit[2]*, G. Zito[1], A. Mangogna[1], Paolo Macor[2], F. Romano[1], M. Toffoli[1], B. Belmonte[3], A. Martorana[4], V. Borelli[2], G. Ricci[1,5], U. Kishore[6] and R. Bulla[2]

**Affiliations**

[1]Institute for Maternal and Child Health, IRCCS Burlo Garofolo, 34137, Trieste, Italy.
[2]Department of Life Sciences, University of Trieste, 34127, Trieste, Italy.
[3]Tumor Immunology Unit, Human Pathology Section, Department of Health Sciences, University of Palermo, Palermo, Italy.
[4]Department of Health Promotion, Mother and Child Care, Internal Medicine and Medical Specialties, University of Palermo, Palermo, Italy.
[5]Department of Medical, Surgical and Health Science, University of Trieste, Trieste, Italy.
[6]Biosciences, College of Health, Medicine and Life Sciences, Brunel University London, London, United Kingdom.

**Supplemental Material**

**Graphical abstract.**

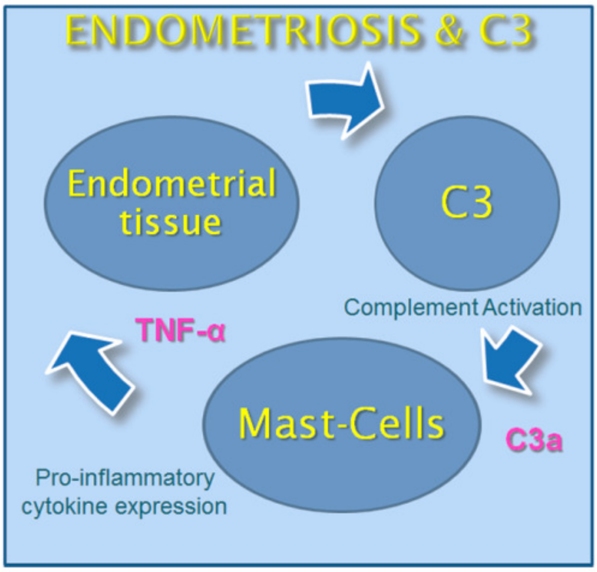

SI Materials and Methods

### Patients for PF Collection

Thirteen women, undergoing either diagnostic or operative laparoscopy at the Institute for Maternal and Child Health, IRCCS Burlo Garofolo, Trieste, Italy were enrolled in a case-controlled study. The study was reviewed and approved by the Regional Ethical Committee of FVG (CEUR), Udine, Italy (Prot. 1197/2015). Informed consent for participation in the study was obtained from all women. The study group (EM) consisted of a total of 7 women, diagnosed with moderate/severe and minimal/mild EM, according to the revised criteria of the American Society for Reproductive Medicine ASRM (*43*). The control group (CTRL) consisted of a total of 6 fertile women, without endometriosis, subjected to laparoscopy to remove leiomyoma. Laparoscopy was performed, and all obtainable peritoneal fluid in the pouch of Douglas was aspirated immediately after entering the abdominal cavity and collected in sterile plastic tubes.

### Cell Lines

HepG2 and AN3CA cells were obtained from American Type Culture Collection (ATCC). HMC-1 cells were kindly provided by Prof. Pucillo (University of Udine, Italy). AN3CA and HepG2 cells were cultured in DMEM/F12, HMC-1 in RPMI-1640, both supplemented with 10% v/v FBS. The medium always contained 100 U/mL penicillin and 100 µg/mL of streptomycin.

### Primary cell isolation and culture

Endometriotic cells and endothelial microvascular uterine cells (UtMEC) were isolated, as described earlier (*44*). ECs were positively selected with Dynabeads M-450 (Life Technologies, Milan, Italy) coated with Ulex europaeus 1 lectin (Sigma-Aldrich), seeded on 12,5 $cm^2$ flask precoated with 2 $\mu g/cm^2$ fibronectin (Roche, Milan, Italy). Cells were maintained in serum-free endothelial basal medium (Life Technologies, Monza, Italy), supplemented with 20 ng/mL bFGF (basic Fibroblast Growth Factor), 10 ng/mL EGF (Epidermal Growth Factor), 10% v/v FBS (all from Life Technologies), and 10% v/v heat inactivated human serum and incubated at 37°C, 5% $CO_2$. Endometriotic epithelial/stromal cells, obtained via the negative selection, were cultured in DMEM containing 10% FBS.

### Murine model of EM

C57BL/6 WT mice were purchased from Harlan Laboratories. $C3^{-/-}$ mice were kindly provided by Prof. Marina Botto, Centre for Complement and Inflammation Research, Department of Medicine, Imperial College, London (UK) and generated as described previously (*45*). All animals were handled in accordance with the institutional guidelines and in compliance with the European (86/609/EEC) and the Italian (D.L.116/92) laws. The Institutional Animal Care Committee of the University of Trieste approved the procedures. The mouse model of EM was adapted from Somigliana and Mariani (*46, 47*). Briefly, donor mice were injected with 17-$\beta$-estradiol-3-benzoate (Sigma-Aldrich; 100 µg/Kg i.m.) and sacrificed 1 week later; the uterus was removed, the two horns were isolated, the myometrium was removed by scraping and the remaining endometrial tissue was reduced to small fragments with scissors. The fragments derived from the isolated uterine horns were weighed and suspended in 400 μl saline (1 mg/ml); half of the preparation was injected into the peritoneum of each of two recipient mice with a syringe (Day 0). Carprofen (5 mg/kg body weight) was given as an analgesic immediately after the surgery and again after 48 h post-surgery. Hormonal therapy with 17-$\beta$-estradiol-3-benzoate (Sigma-Aldrich, 50 μg/kg i.m.) was initiated at the time of tissue injection and at 2-day interval thereafter. Mice were sacrificed at day 21 via administration of a lethal dose of anesthetic, their abdomen was opened, and the presence of the lesions was evaluated by an operator blinded to the different conditions. Translucid isolated or grouped superficial lesions were mainly found on the abdominal wall, on the epiploon and around the uterus. Deeply infiltrating lesions were never observed in this model. In some cases, lesions resembling human chocolate cysts were found.

**Characterization of EM cells**

EM cells were plated on 8-chamber culture slides (BD Biosciences Discovery Labware, Milan, Italy). Cells, when grown to confluence, were fixed and permeabilized with FIX & PERM (Società Italiana Chimici, Rome, Italy). Next, cells were incubated with primary monoclonal antibody (mAb) (clone 9) mouse anti-human vimentin (Sigma-Aldrich), (cloneF8/86) mouse anti-human vWF (Dako-Cytomation, Milan, Italy), or mouse anti-human CK 8/18 (Abcam, Italia, Milan, Italy) for 1 h at room temperature (RT), followed by fluorescein isothiocyanate (FITC)-conjugated goat anti-mouse IgG for 1 h at RT. Images were acquired using Leica DM3000 microscope (Leica, Wetzlar, Germany) and collated using a Leica DFC320 digital camera (Leica).

**Western Blot Analysis**

$10^6$ ANC3CA cells seeded onto 6-well plates were treated with 100 ng/mL of TNF-α or 5 ng/mL of IL-1β for 36 h. Cells lysates were fractioned by 10 % SDS-PAGE under reducing conditions and transferred to a nitrocellulose membrane using the semi-dry transfer apparatus Trans-Blot Turbo System according to the manufacturer's protocol (BIO-RAD). 100 ng of recombinant human C3 (Quidel) were used as a control.

After 1 h of incubation with 5 % skim milk in TBST (10 mM Tris, pH 8.0, 150 mM NaCl, 0.5% Tween 20), the membrane was probed with 1:500 anti-C3 (MyBioSource) ON at 4°C. Membrane was washed three times for 5 min and incubated with 1:10000 anti-goat LI-COR IRDye 800CW for 1h at RT. After three washing steps, the fluorescence intensity was acquired by the Odyssey® CLx near-infrared scanner (LI-COR Biosciences, Lincoln, NE, USA). Image acquisition, processing and data analysis were performed with Image Studio Ver 5.2 (LI-COR Biosciences).

**Immunohistochemical analysis**

Uterine and ectopic endometrial tissues were enrolled in this study, after approval of the University Hospital of Palermo Ethical Review Board (approval number 09/2018). Our study selected two cases of proliferative and secretory endometrium as controls and two cases of patients with tubal and abdominal wall endometriosis.

Immunohistochemistry (IHC) was performed using a polymer detection method. Briefly, tissue samples were fixed in 10% v/v buffered formalin and then paraffin embedded. 4 μm-thick tissue sections were deparaffinized and rehydrated. The antigen unmasking technique was performed using Novocastra Epitope Retrieval Solutions, pH 6 EDTA-based (Leica Biosystems) in thermostatic bath at 98°C for 30 min. Sections were then brought to RT and washed in PBS. After neutralization of the endogenous peroxidase with 3% v/v $H_2O_2$ and Fc blocking by a specific protein block (Novocastra, Leica Biosystems), samples were incubated for 1 hour at RT with mouse anti-human C3a/C3a (dilution 1:50 ph6) monoclonal antibody (Millipore) and rabbit anti-human C3 (dilution 1:200 ph9) polyclonal antibody (Sigma-Merk)). Staining was revealed via polymer detection kit (Novocastra, Leica Biosystems) and AEC (3-amino-9-ethylcarbazole, Dako, Denmark) substrate chromogen. Slides were counterstained with Harris Hematoxylin (Novocastra, Leica Biosystems). Toluidine blue stain was carried out to detect the presence of mast cells in uterine and ectopic endometria tissues, according to the manufacturer's kit Histoline.

Slides were analyzed under the Axio Scope Sections were analyzed under the Axio Scope A1 optical microscope (Zeiss) and microphotographs were collected through the Axiocam 503 color digital camera (Zeiss) using the Zen2 software.

### Tryptase Concentration and Enzymatic Activity

Tryptase ELISA kit (USCN, Life Sciences Inc) was used to determine the concentration of tryptase in mice PF samples. The assays were performed according to the manufacturer's instructions and the results referred to a calibration curve expressed in ng/mL. Samples were assayed in triplicate.

### Statistical Analysis

Data were analyzed by GraphPad Prism software 5.0 (GraphPad Software Inc., La Jolla, CA, USA). Unpaired two-tailed Mann-Whitney test was used for the analysis of different tissue C3 gene expression and cells; Wilcoxon test was applied for AN3CA stimulation and mouse model. Results were expressed as mean ± SEM of three independent experiments performed in duplicate. P-values $<0.05$ were considered statistically significant.

### Supplementary Materials

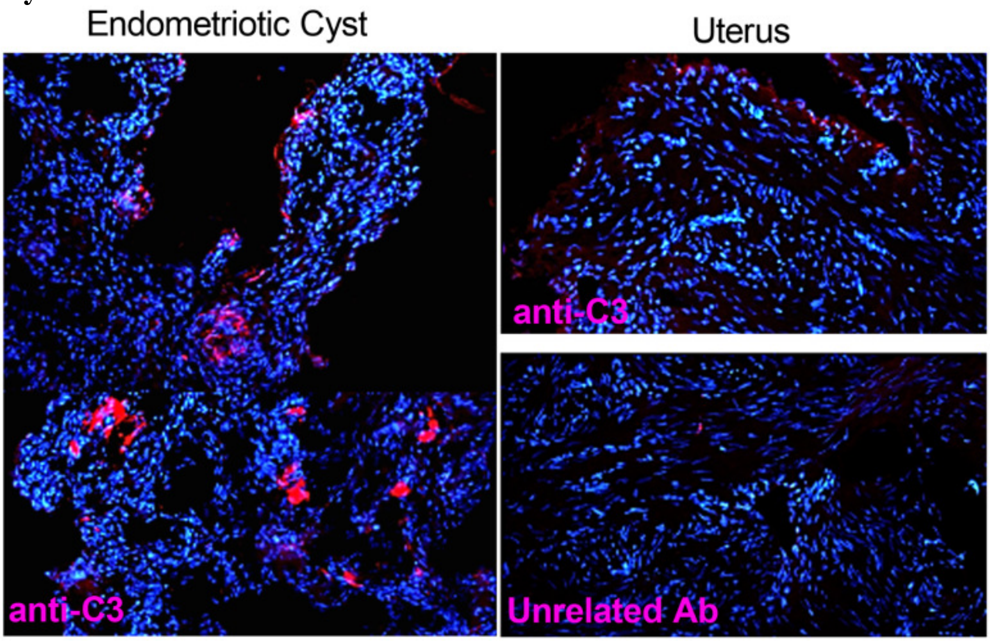


**Fig. S1.** Immunofluorescence analysis of endometrial eutopic and ectopic tissue on frozen sections for C3 presence. EM cyst (left part) and uterine endometrium (right part). C3 staining (with goat anti-human C3, Quidel, Rome, Italy), in red, was strongly marked in endometriotic cyst section compared to normal uterus. The nuclei were contra-stained in blue with DAPI. (original magnification 100X).

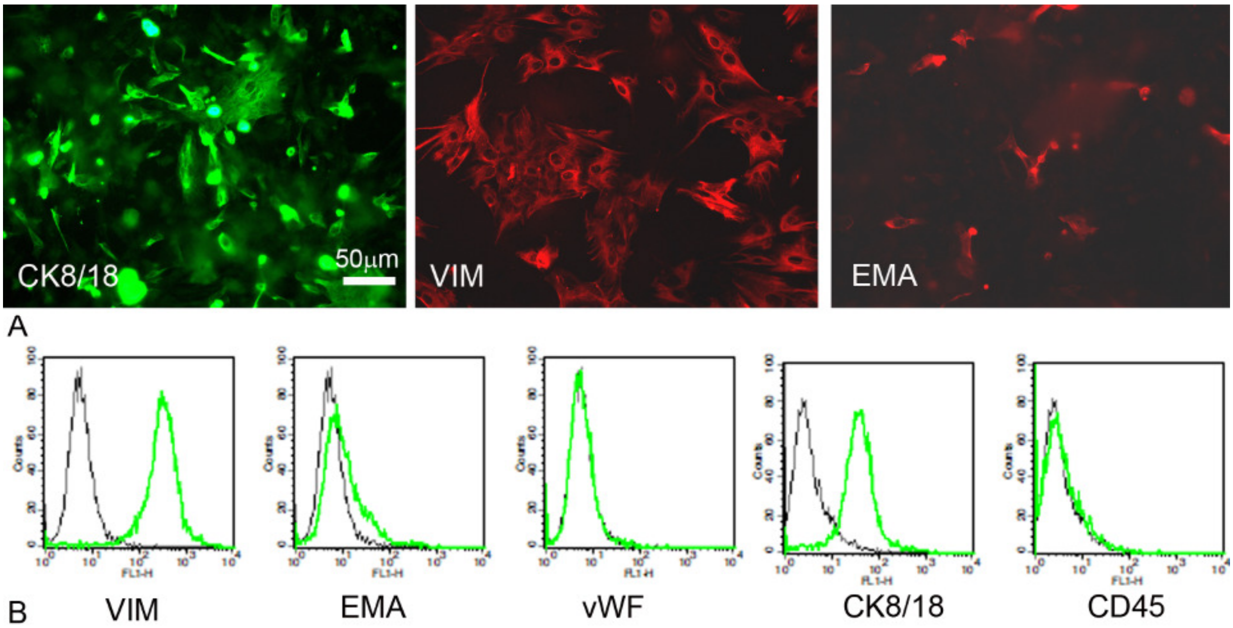


**Fig. S2.** (A) Endometriotic cells were characterized by immunofluorescence for the expression of cytokeratin 8/18 (CK) in green, vimentin (VIM) and Epithelial Membrane Antigen (EMA) in red. The cells were grown to confluence in eight-chamber culture slides. After fixation and permeabilization, the cells were stained with mAb primary followed by anti-mouse-FITC or –Cy3 conjugated F(ab)′ secondary antibodies. Original magnification 200×. (B) The expression of Vimentin, EMA, von Willebrand Factor, CK8/18, and CD45 was analyzed by cytoflurimetric analysis. The expression of these markers (green lines) was compared with appropriate control antibodies (black lines).

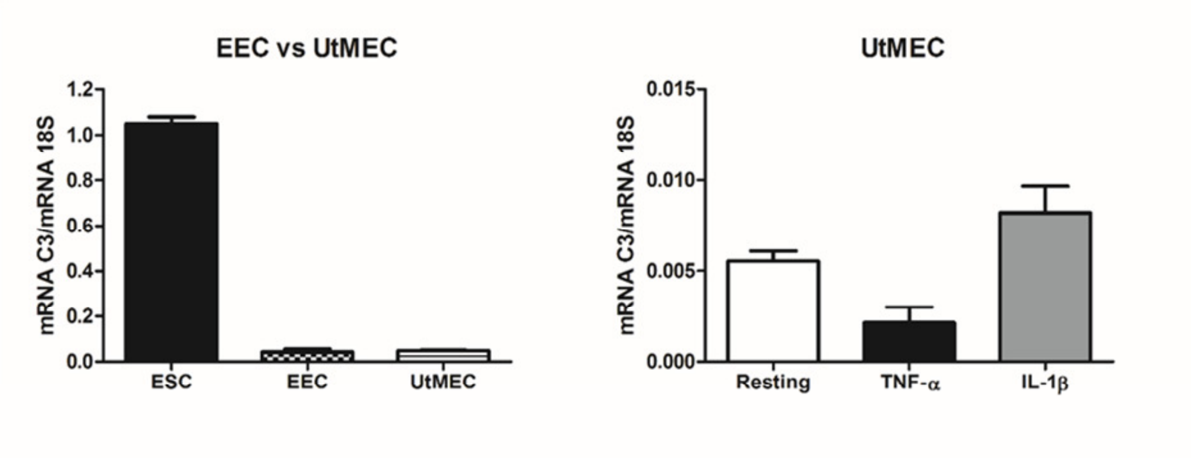


**Fig. S3.** (A) The C3 mRNA expression was evaluated by RT-qPCR in Endometriotic Endothelial Cells (EEC, n = 4) and compared to an Uterine Microvascular Endothelial Cells (UtMEC, n = 3). Endometrial cell isolated from EM cyst (ESC) were used as calibrator. (B) UtMEC were O.N. stimulated with TNF-α (100 ng/mL) or IL-1β (5 ng/mL) and the C3 expression was analyzed by RT-qPCR. No significant modulation of C3 expression was observed. Data are expressed as mean ± standard error.

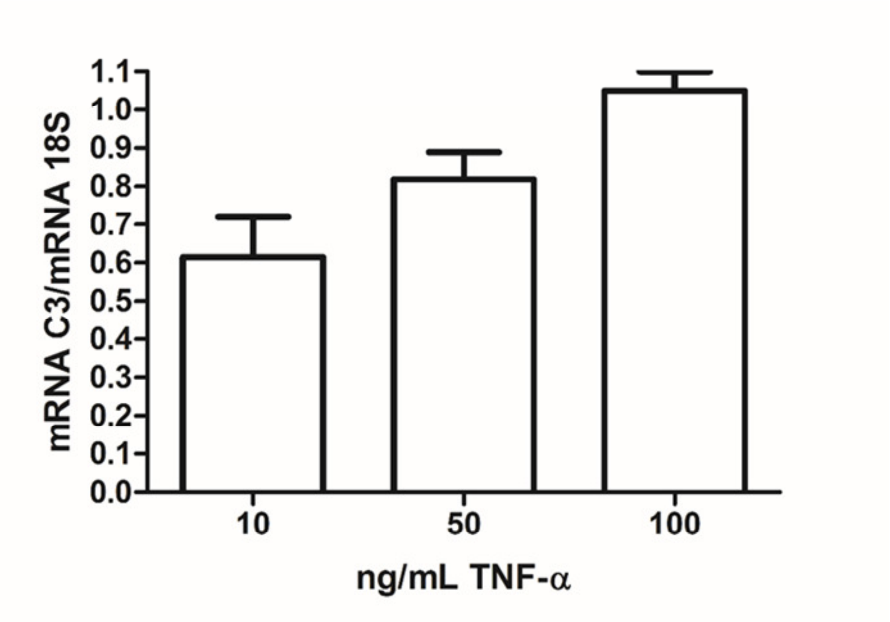


**Fig. S4.** TNF-α induced expression of C3 mRNA by AN3CA in a dose dependent manner. The C3 mRNA expression was evaluated by RT-qPCR in AN3CA after 24h stimulation of increasing concentrations of TNF-α (10, 50, 100 ng/mL). Data are expressed as mean ± standard error.

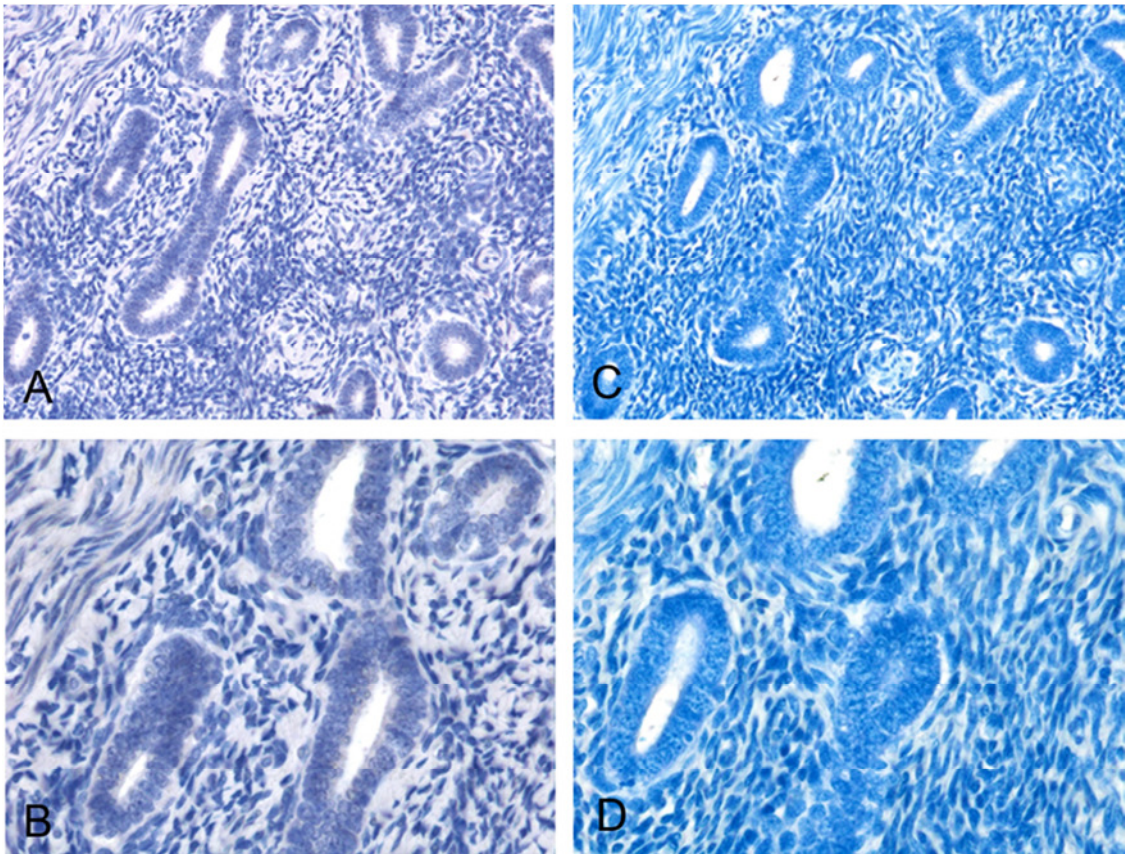


**Fig. S5.** IHC and HC analysis of C3a (A and B) and mast cells (C and D) presence respectively, in secretive uterine endometrium. IHC analysis and toluidine blue staining revealed the absence of both C3a and mast cells in secretory endometrium.

| Gene | Tm (°C) | Sense | Sequence (5′-> 3′) | Accession number |
|---|---|---|---|---|
| **18 S** | 61 | Forward | ATCCCTGAAAAGTTCCAGCA | NM_022551.2 |
| | | Reverse | CCCTCTTGGTGAGGTCAATG | |
| **C3** | 61 | Forward | CCTGCTACTAACCCACCTCC | NM_4557385a1 |
| | | Reverse | AACAGTGACTGGAACATCCCC | |
| **C5** | 60 | Forward | ATGGGCCTTTTGGGAATACTTTG | NM_4502507a1 |
| | | Reverse | ACATGGCCTGAGGAGTAACTAA | |

**Table S1.** Primer used for RT-qPCR analysis.
